# Copy number variations in mitochondrial DNA has a potent role in parasitic immunity against *Haemonchus contortus-*A novel report

**DOI:** 10.1101/2025.09.10.675310

**Authors:** Aruna Pal, Samiddha Banerjee, Kavita Rawat, Debapritam Deb, P.N. Chatterjee

**Author notes:** Corresponding author: E mail.

## Abstract

Mitochondria is a cellular organelle responsible for a variety of function, apart from energy production through oxidative phosphorylation, TCA cycle, apoptosis, plays a major role in iron and calcium metabolism and immune functions against pathogens (bacterial, viral or parasitic). Unlike nucleus, mitochondria exist in multiple copies, with many copies of mitochondrial DNA. A metabolically active cell possess about 10^3^ and 10^4^ copies of the mitochondrial genome. Lowered mtDNA copy number leads to disorders or disease, in a similar way as mutation in mtDNA causes a wide range of disease or disorders. In this current article, we estimated mtDNA copy number through QPCR analysis of cytochrome B gene and revealed that healthy individuals have better mtDNA copy number compared to that of the infected ones under natural challenge in sheep model against the parasite *Haemonchus contortus* infection. We diagnosed the infected animals for *H. contortus* primarily with symptoms, followed by Faecal egg count and final confirmation through molecular diagnosis. We had characterized the cytochrome B gene from abomasum(stomach) of sheep, which is the site of predilection of this parasite. In our lab, earlier we have documented certain immune response nuclear genes to have antiparasitic role. We had reported the mutations in cytochrome B leading to debility and disease in sheep. Quantitative expression profile for cytochrome B is an indication of the copy number of mitochondrial genes and we observed significantly better expression profile of cytochrome B in healthy sheep in comparison to infected one. Thus, mitochondrial copy number is directly associated with antiparasitic immunity, first time reported.

## Introduction

Mitochondria and its DNA content has quite unique features and have distinct features than that of nucleus and nuclear DNA. There exists certain interesting facts about mitochondria. Mitochondrial division occurs through simple fission, splitting in two like bacterial cells. However the DNA replication strategies differs slightly, forming displacement or D-loop structures and they partition their circular DNA similar to bacteria. Mitochondria is considered as autonomous cell organelle due to the certain reasons. Mitochondria have their own DNA which can replicate independently^1–3^. It is interesting to note that most of the mitochondrial proteins are synthesized under genetic control of cell nucleus. Mitochondria can thus be described as double walled semi-autonomous, contains DNA, an oxidative and energy transducing organelle. It is found only in Eukaryotic aerobic cells (*except mature RBCs totally absent in prokaryotes & anaerobic eukaryotic cells).* Mitochondria is semiautonomous because <u>i</u>t contain their own DNA, have cytochromes ETS for ATP synthesis, mitochondria is arise from pre existing ones from division i.e. *not de novo, have* DNA forming protein used in making their own membranes. But enzymes in mitochondria are coded by nuclear DNA (*therefore mitochondria is not fully autonomous it is semiautonomous)* ^1–3^. Three most important nuclear coded genes affecting Mitochondrial function DNA polymerase, Polymerase Gamma (POLG), transcription factor TFAM and DNA helicase as TWINKLE^4^. Mitochondrial copy number is greatly influenced by important mitochondrial phenomenon as mitochondrial fusion, mitochondrial fisson and mitophagy, they are in turn controlled by nuclear coded genes as mitofusin and others. Mitochondria have their own DNA which can replicate independently^5,6^. The *mitochondrial DNA* synthesizes its own *mRNA, tRNA and rRNA.* The organelles posses their own ribosomes, called *mitoribosomes*^6–8^. Mitochondria synthesize some of their own structural proteins. However, most of the *mitochondrial proteins are synthesized under instructions from cell nucleus.* The organelles synthesize some of the enzymes required for their functioning. *e.g. succinate dehydrogenase*. They show hypertrophy. *i.e.* internal growth. Existence of nuclear mitochondrial genes (NUMT) further makes the process interesting. Anterograde and retrograde system further make mitochondria and nuclear interaction further complicated and interesting. Mitochondrial protein affect immune system through certain nuclear immune response proteins, namely NRLP3, IL18, STING we studied earlier against bacterial infection^9^.

In our earlier studies, we have studied and characterized Cytochrome B gene in garole sheep. We have already identified certain SNPs pertaining to cytochrome B and identified certain deleterious mutations affecting sheep health and debility^1^. SNPs were identifed through sanger sequencing^10^. Cytochrome B has a multifaceted functions for metal ion binding (iron, calcium), ubiquinol-cytochrome-c reductase activity, and mitochondrial electron transport, ubiquinol to cytochrome C^1,2, 11^.

Assessment of livestock health conditions in developing countries for identification of priority diseases to be targeted for control, revealed helminth infections as one of the most important problems in sheep and goat^11^. Gastro-intestinal parasitic infestations such as *Haemonchus contortus,* impose severe constraints on sheep and goat production especially those reared by marginal farmers under low external input system. These parasites results in heavy losses to farmers due to its body weight loss, direct cost of anthelminthic drugs, loss due to mortality, etc. For example, annual treatment cost for *Haemonchus contortus* alone had been estimated to be 26 million USD in Kenya, 46 million USD in South Africa and 103 million USD in India^12^. Generally sheep were observed to be hardy and better resistant compared to goat^13,14^. Emergence of strains resistant to anthelminthic drugs has further complicated the management of parasitic diseases in small ruminants^15–16^. Genetic variation in host resistance exists for the major nematode species affecting sheep: *Haemonchus contortus*, Considerable variation has been reported among sheep breeds on their ability to resist gastro-intestinal nematodes (GIN)^11^. In our lab, we have already detected certain nuclear immune response genes responsible for antiparasitic immunity, as RIGI^17^, CD14^18^, MyD88^19^, IL6^20^, IL10^20^. We have equally studied whole mitochondrial genome sequencing for sheep^21^ and duck^22,23^.

Considering the facts, in this current study, we aim to detect if there is any role for mitochondrial copy number with antiparasitic resistance. We have studied sheep as the model organism and *Haemonchus contortus* as the infective agent.

## Materials and Method

### Animals

We randomly collected 60 Garole sheep (*Ovis aries*) faecal samples from the Livestock Farm, West Bengal University of Animal and Fishery Sciences, Mohanpur campus. The samples were collected prior to a routine deworming procedure and were presumed to have preexisting gastrointestinal parasites because they were allowed to graze freely. The animals were exposed to natural infection during grazing.

### Estimation of Faecal egg count

Fecal samples from sheep were examined using the salt flotation method [17–20], and fecal egg counts (FECs) were determined for each sample. Approximately 1 g of fecal material was homogenized with a mortar and pestle, followed by the addition of 15 ml of saturated NaCl solution. After sedimentation, the supernatant was collected with a dropper and transferred into a McMaster counting chamber by capillary action. The preparations were subsequently examined microscopically. FEC was calculated using the formula:

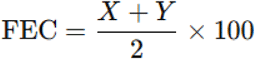

where *X* and *Y* represent the egg counts in the two chambers of the McMaster slide.

For statistical evaluation, animals were classified into two groups: “healthy” (mean + SD) and “diseased” (mean – SD), where mean denotes the arithmetic average and SD represents the standard deviation. Sheep used in the study were part of the regular stock sold and slaughtered for mutton production. In total, twelve abomasal tissue samples were analyzed, comprising six with low FEC values (designated as healthy) and six with high FEC values (classified as infected). The presence of infection was further confirmed through direct visualization of adult helminths in the abomasum.

Tissue samples were collected from the abomasum.

### Haematological Profiles

The haematological parameters like haemoglobin, erythrocyte sedimentation rate (ESR) and packed cell volume (PCV) were assessed from whole unclotted blood. Haemoglobin was estimated by acid haematin method^18,20,24^, E.S.R. and PCV by Wintrobe’s tube^24^. The total erythrocyte count (TEC), total leukocyte count (TLC) and Differential leukocyte count (DLC) were studied by standard methods described by Jain^18,20,26^,. Biochemical Analysis The serum biochemical parameters estimated in the experiments were total protein, albumin, globulin, albumin: globulin, aspartate aminotransferase (AST), alanine aminotransferase (ALT), alkaline phosphatase (ALP), Total bilirubin, Indirect bilirubin, direct bilirubin, glucose, uric acid, urea, and BUN by using a semi auto biochemistry analyser (Span diagnostic Ltd.) with standard kits (Trans Asia Bio-Medicals Ltd., Solan, HP, India). The methodology used for estimation of total protein, albumin, total & direct bilirubin, ALT, ALP, glucose, creatinine urea, and uric acid were biuret method, bromocresol green (BCG)method,2-4 DNPH method, modified kind and king’s method, GOD/POD method, modified Jaffe’s Kinetic method. GLDH-urease method, and trinder peroxidise method respectively^18,20^.

### Molecular detection and confirmation of Haemonchus contortus from faeces

#### Characterization of Cox1 gene of *Haemonchus contortus*

Salt floatation method was employed for concentrating the faecal eggs. DNA isolation was followed DNA with standard Phenol chloroform isoamyl alcohol process^27^ for hatched embryos also.

#### Molecular diagnosis for Haemonchus contortus and quantification through QPCR

Genomic DNA was isolated from faeces sample from both parasitic eggs as well as hatched larvae. PCR amplification for Cox1 gene of *Haemonchus contortus* was performed. Positive samples with CT values confirm the detection of *Haemonchus contortus*.

#### Real-Time PCR Experiment (SYBR Green based)

The primers were standardised with respective cDNA samples before being subjected to real-time PCR. The reactions were conducted in triplicate (as per MIQE Guidelines) with the total volume of the reaction mixture made up to 20µl. The experiment was conducted with Cytochrome B gene and GAPDH (housekeeping gene) primers calculated as per concentration. 1µl of cDNA was mixed with 10µl Hi-SYBr Master Mix (HIMEDIA MBT074) and volume adjusted upto 20µl with nuclease-free water. Then the reaction was performed into the ABI 7500 system and run the reaction program. The delta-delta-Ct (ΔΔCt) method 2^-ΔΔCt^ was used for the analysis of the expression level od particular gene^27–31^. The primers used for the reaction are as followed:

### Characterization of cytochrome B gene from abomasum of sheep mRNA Isolation

Gut tissue (abomassum) from healthy and diseased sheep were collected followed by total RNA isolation by the TRIzol method to assess gut-associated lymphoid tissue (GALT). Abomasal tissues were collected individually from both healthy and *Haemonchus contortus*– infected sheep. Total RNA was extracted under aseptic conditions using the TRIzol reagent method. For each animal, abomasal tissue samples were systematically labeled. mRNA was withdrawn from abomasum tissue by the TRIzol method and subjected to cDNA preparation^27–31^. Each mixture of 2 g of tissue and 5 ml TRIzol was triturated, followed by chloroform treatment. Centrifugation resulted in three-phase differentiation from which the aqueous phase containing the mRNA was separated. It was treated by isopropanol to extract the mRNA in the form of a pellet, discarding the supernatant as a result of centrifugation. Finally, the pellet underwent an ethanol wash and was air-dried, followed by dissolving the mRNA in nuclease-free water.. RNA concentration and quality was estimated by nanodrop as per standard procedure^27–31^.

#### Materials Required

10X buffer, dNTP, and Taq DNA polymerase were procured from Invitrogen; SYBR Green qPCR Master Mix (2X) was obtained from Thermo Fisher Scientific Inc. (PA, USA). Primers were purchased from Xcelris Labs Limited. The reagents used were of analytical grade.

### Configuration of cDNA, and PCR Amplification of cytochrome B Gene

First, 20 μL reaction mixture volume consisted of 5 μg of complete RNA, 40LU of ribonuclease inhibitor, 0.5 μg of oligo dT primer (16–18Lmer), 1000LM of dNTP, 5LU of MuMLV reverse transcriptase in reverse transcriptase buffer, and 10LmM of DTT. The reaction mixture was carefully blended and incubated properly at 37°C for 1 h. The reaction was paused and heated at 70°C for 10 min followed by immediate chilling on ice. The quality of the cDNA was assessed by nano drop and gel electrophoresis and by polymerase chain reaction.

25μL of reaction mixture was comprised of 80–100ng cDNA, 3.0μL 10X PCR assay buffer, 0.5μL of 10mM dNTP, 1U Taq DNA polymerase, 60ng of each primer, and 2mM MgCl2. PCR reactions were conducted in a thermocycler (PTC-200, MJ Research, USA), cycling conditions being initial denaturation at 94°C for 3min, further denaturation at 94°C for 30sec, annealing at 61°C for 35sec, and extension at 72°C for 3 min were conducted for 35 cycles followed by a final extension at 72°C for 10 min. The list of primers pertaining to these genes are listed in Table 2:

**Table 1:**
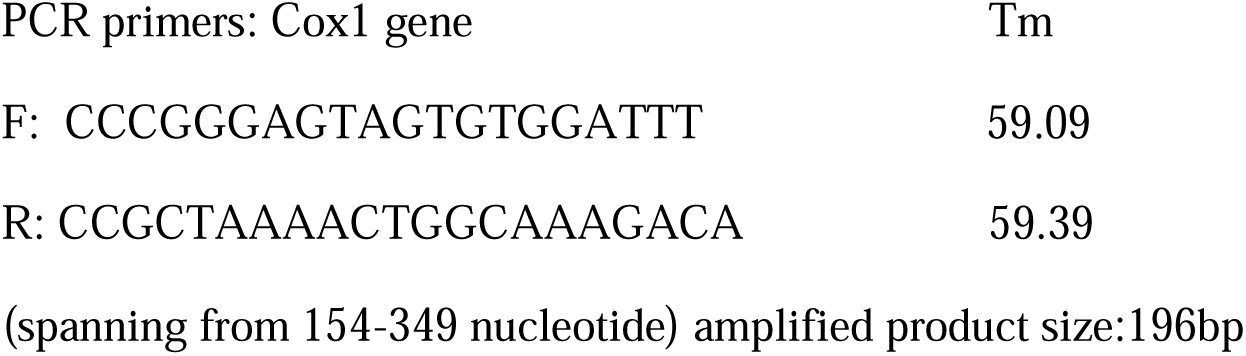
PCR primers for mitochondrial cox1 gene from *Haemonchus contortus*.

**Table 2:**
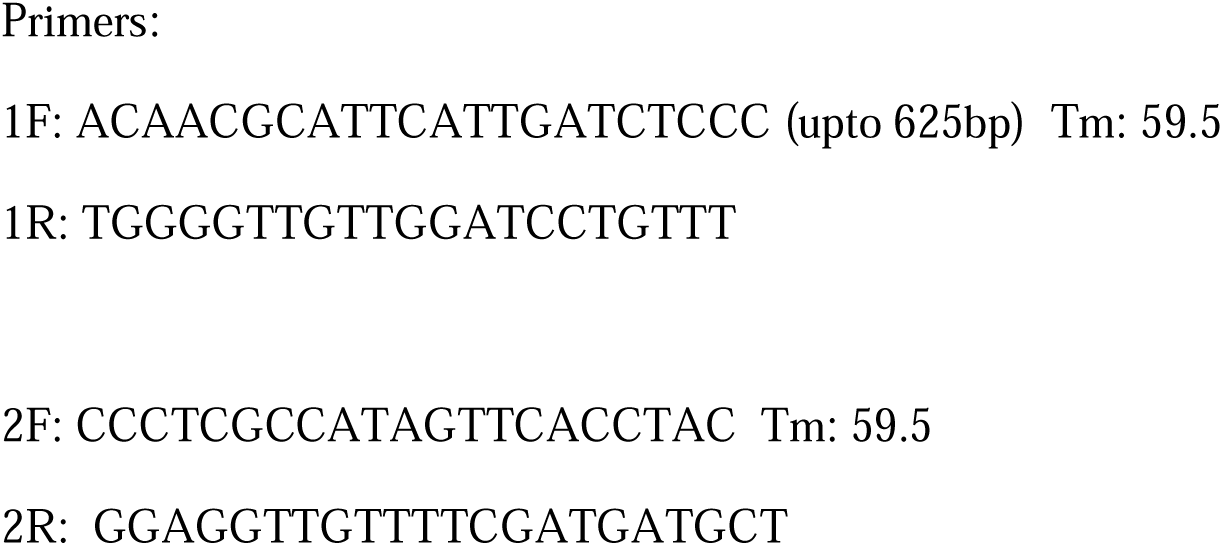
PCR primers for cytochrome B gene from abomasum of sheep Primers:

To obtain a full-length open reading frame (ORF) of the gene sequence, specific primers pairs were designated for the amplification of cytochrome B and confirmed based on the mRNA sequences of *Bos indicus* by DNASTAR software. The amplified products were designed in such way to have the overlapping sequences that can be joined to extract the full sequence.

### Study of Mitochondrial Cytochrome B of sheep *in silico*

Predicted peptide sequences of duck **Mitochondrial Cytochrome B we**re analysed. The analysis was conducted for a sequence-based comparative study. The signal peptide is essential to prompt a cell to translocate the protein, usually to the cellular membrane and ultimately signal peptide is cleaved to give a mature protein. Prediction of the presence and location of the signal peptide of these genes was conducted using the software (Signal P 3.0 Sewer-prediction results, Technical University of Denmark). The leucine percentage was calculated manually from the predicted peptide sequence. Di-sulphide bonds are necessary for protein folding and stability. It is the 3D structure of the protein which is biologically active.

Protein sequence level analysis was performed (http://www.expasy.org./tools/blast/) for the assessment of leucine-rich repeats (LRR), leucine zipper, estimation of Leucine-rich nuclear export signals (NES), and prediction of the position of GPI anchor, N-linked glycosylation sites. Being receptors, they are rich in leucine-rich repeats, necessary for pathogen recognition and binding. A leucine zipper is essential to assess the dimerization of IR molecules. N-linked glycosylation is important for the molecule to assess its membranous or soluble form. The leucine-rich nuclear export signal is essential for the export of this protein from the nucleus to the cytoplasm, whereas GPI anchor is responsible for anchoring in the case of membranous protein. Prosite was employed for LRR site detection.

Leucine-rich nuclear export signals (NES) were analyzed with NetNES 1.1 Server, Technical University of Denmark. O-linked glycosylation sites were assesed using NetOGlyc 3.1 server (http://www.expassy.org/), and N-linked glycosylation sites were predicted through NetNGlyc 1.0 software (http://www.expassy.org/). Sites for leucine-zipper were detected through Expassy software, Technical University of Denmark^32^. Sites for alpha helix and beta sheet were detected using NetSurfP-Protein Surface Accessibility and Secondary Structure Predictions, Technical University of Denmark^33^. Domain linker sites were predicted^34^. LPS-binding^35^ and LPS-signalling sites^36^ were predicted based on homology studies with other species of respective polypeptide. These sites are important for pathogen recognition and binding.

### Three-dimensional structure prediction and Model quality assessment

3D models of **Mitochondrial protein (Cytochrome B & Cytochrome C) and nuclear protein (NRLP3, IL18 and Sting)** polypeptide were modelled through the Swissmodel repository^37^. Templates containing the greatest identity of sequences with our target template were identified with PSI-BLAST (http://blast.ncbi.nlm.nih.gov/Blast). PHYRE2 server based on the ‘Homology modelling approach was used to build three dimensional model of these proteins^38^. Molecular visualization tool as PyMOL (http://www.pymol.org/) was employed for model generation and visualization of the three-dimensional structure of these proteins understudies for duck origin. The structure of duck molecules was evaluated and assessed for its stereochemical quality (through SAVES, Structural Analysis and Verification Server, http://nihserver.mbi.ucla.edu/SAVES/); then refined and validated through ProSA (Protein Structure Analysis) web server (https://prosa.services.came.sbg.ac.at/prosa)^39^. NetSurfP server (http://www.cbs.dtu.dk/services/NetSurfP) was employed for assessing the surface area of these proteins with relative surface accessibility, Z-fit score, and probability of alpha-Helix, beta-strand and coil score.

The alignment of 3-D structure of these proteins was checked with RMSD estimation to assess the structural differentiation by TM-align software^40^.

### Protein-protein interaction network depiction

To understand the protein interaction network of these proteins, we performed the search in STRING 9.1 database^41^. The functional interaction was assesed with a confidence score. Interactions with scores < 0.3, scores ranging from 0.3 to 0.7, and scores >0.7 are designated as low, medium and high confidence respectively. We have also conducted a KEGG analysis depicting the functional association of these proteins with other related proteins.

### Differential mRNA expression for Cytochrome B gene for Sheep w.r.t to healthy vs. infected

#### mRNA Isolation

Abomassal tissues from healthy and diseased sheep were collected and followed total RNA isolation by the TRIzol method to assess gut-associated lymphoid tissue (GALT) as we described during Characterization of cytochrome B section. Total RNA was extracted under aseptic conditions using the TRIzol reagent method. For each animal, abomasal tissue samples were systematically labeled. mRNA was withdrawn from abomasum tissue by the TRIzol method and subjected to cDNA preparation^27–31^. Each mixture of 2 g of tissue and 5 ml TRIzol was triturated, followed by chloroform treatment. Centrifugation resulted in three-phase differentiation from which the aqueous phase containing the mRNA was separated. It was treated by isopropanol to extract the mRNA in the form of a pellet, discarding the supernatant as a result of centrifugation. Finally, the pellet underwent an ethanol wash and was air-dried, followed by dissolving the mRNA in nuclease-free water.. RNA concentration and quality was estimated by nanodrop as per standard procedure^27–31^.

#### Materials Required

10X buffer, dNTP, and Taq DNA polymerase were procured from Invitrogen; SYBR Green qPCR Master Mix (2X) was obtained from Thermo Fisher Scientific Inc. (PA, USA). Primers were purchased from Xcelris Labs Limited. The reagents used were of analytical grade.

### Configuration of cDNA, and PCR Amplification of cytochrome B Gene

First, 20 μL reaction mixture volume consisted of 5 μg of complete RNA, 40LU of ribonuclease inhibitor, 0.5 μg of oligo dT primer (16–18Lmer), 1000 M of dNTP, 5LU of MuMLV reverse transcriptase in reverse transcriptase buffer, and 10 mM of DTT. The reaction mixture was carefully blended and incubated properly at 37°C for 1 h. The reaction was interrupted and heated at 70°C for 10 min, next immediately chilled on ice. The quality of the cDNA was checked by nanodrop OD and polymerase chain reaction. To get a full-length open reading frame (ORF) of the gene sequence, the primers pairs were assigned for the amplification of Cytochrome B and confirmed based on the mRNA sequences of *Bos indicus* by DNASTAR software as in <u>Table</u>. The amplified products containg the overlapping sequences joined to extract the full sequence.

### Real-Time PCR Experiment (SYBR Green based)

The primers were standardised with respective cDNA samples before being subjected to real-time PCR. The reactions were conducted in triplicate (as per MIQE Guidelines) with the total volume of the reaction mixture made up to 20µl. The experiment was conducted with Cytochrome B and GAPDH (housekeeping gene) primers calculated as per concentration. 1µl of cDNA was added to 10µl Hi-SYBr Master Mix (HIMEDIA MBT074) and rest volume adjusted upto 20µl with nuclease-free water. Then the reaction was conducted in the ABI 7500 system. The delta-delta-Ct (ΔΔCt) method 2^-ΔΔCt^ was used for the analysis of the result^27–31^. The primers used for the reaction are as followed:

### Statistical analysis

ANOVA was employed for the analysis of haematological and biochemical analysis of the infected sheep with the healthy ones. Each sample was run in triplicate. Analysis of real time PCR (qRT-PCR) was performed by delta-delta-Ct (DDCt) method, Ct denotes the threshold value.

## Result

### Characterization of cytochrome B gene from sheep abomasum

Sequencing of cytochrome B gene from gut tissue (abomassal tissue) of sheep was characterized (gene bank accession number **2996374**.). 3D structure was predicted with important domains identified (Fig 1), revealing four sites for iron binding and one ubiquinone site. Transmembrane helical domains for cytochrome B have been identified as revealed in Fig 2. Cytochrome B works in conjunction with multiple other proteins, majority of them are cytoplasmic while some nuclear. Molecular interaction of cytochrome B with other proteins are being depicted through String analysis (Fig 3). One of the major role for cytochrome B is in immunity against infections, that they execute through multiple mechanisms as defect in phagosome pathway (Fig 4) and hindrance in biosynthesis of various nucleotide sugar (Fig 5), lead the animal more prone to diseases.

**Fig 1:**
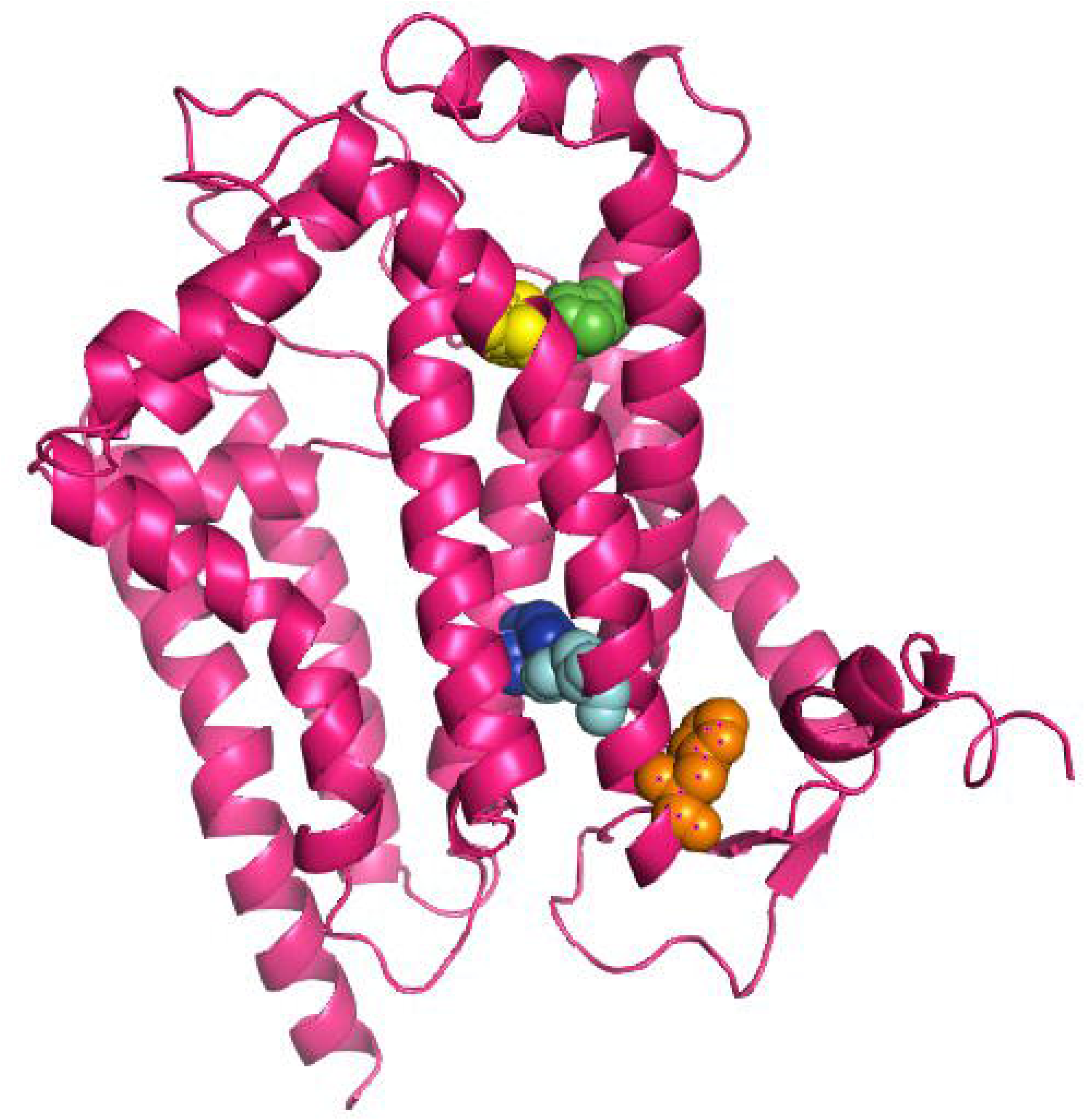
3D structure of cytochrome B gene from abomassum of sheep (hot pink) Binding sites for iron at amino acid positions 83(green sphere), 97(blue sphere), 182(yellow sphere), 196(cyan sphere), and ubiquinone binding site at a.a position 201 (orange sphere).

**Fig 2:**
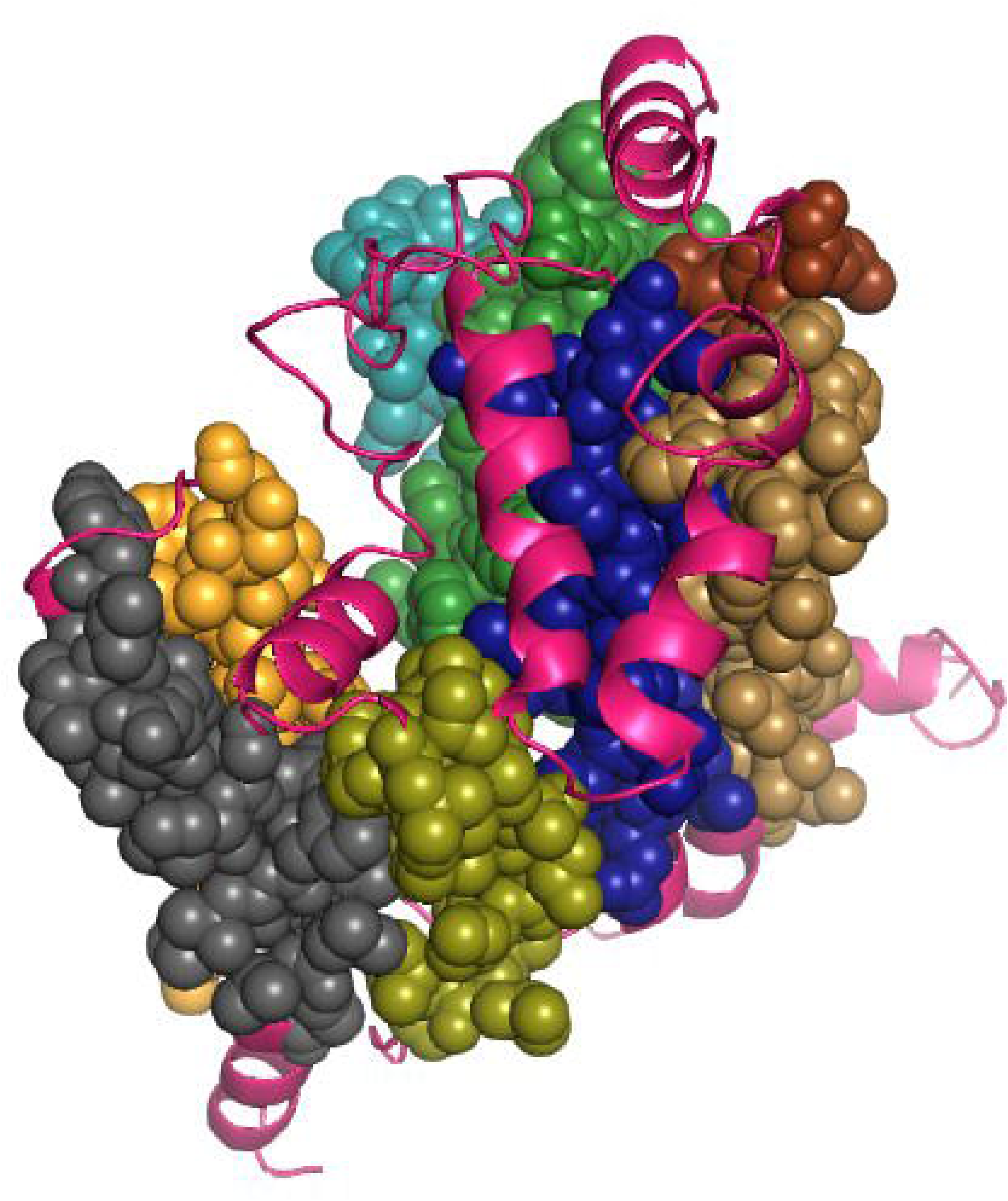
3D structure of cytochrome B gene from abomasum of sheep (hot pink) with transmembrane helical sites at amino acid positions 33-53 (chocolate sphere), 77-98(forest green sphere), 113-133(density blue sphere), 178-198(sand yellow sphere), 226-246(deep teal sphere), 288-308 (deep olive sphere), 320-340 (bright orange sphere), 347-367(grey sphere).

**Fig 3:**
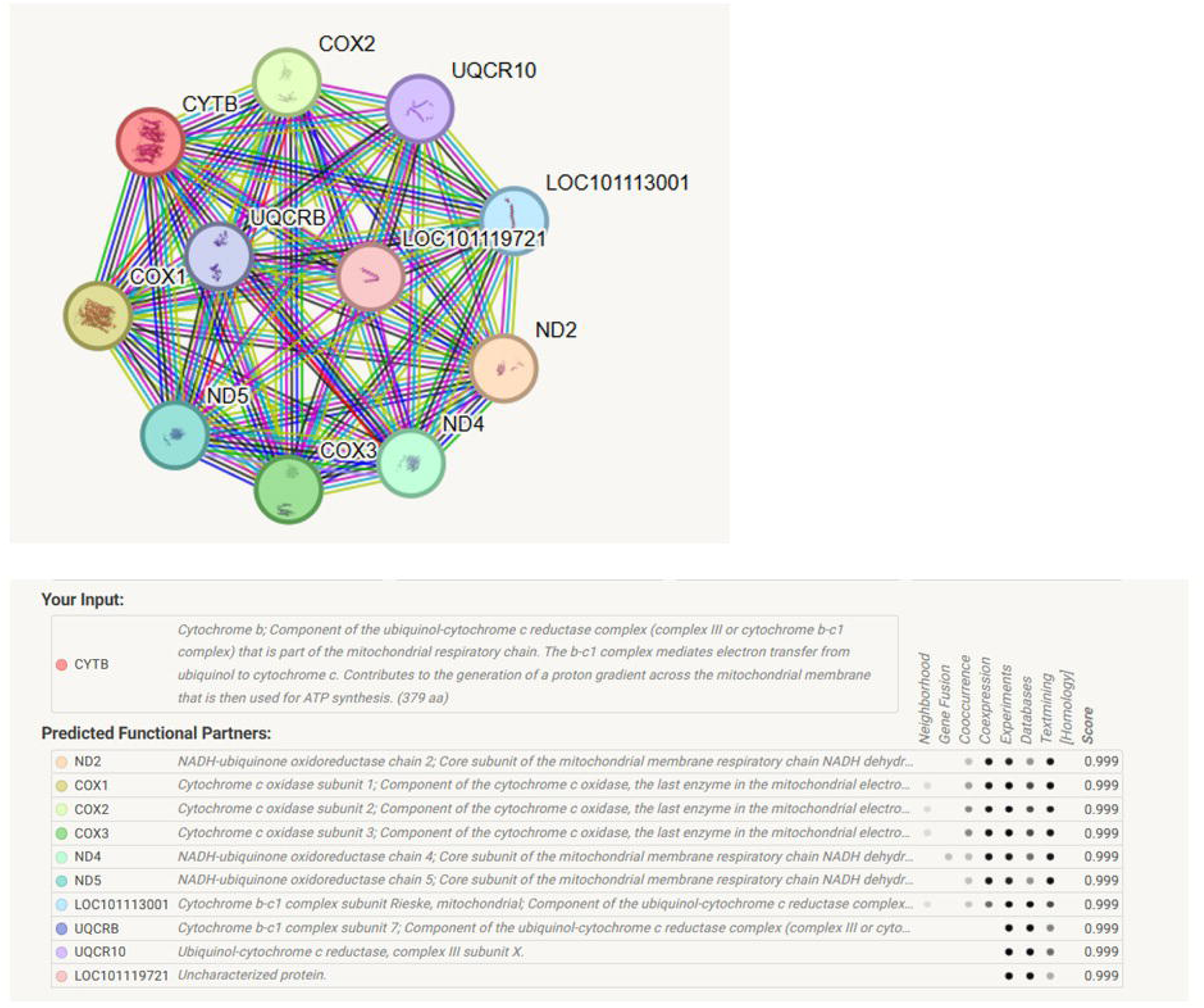
String Analysis for molecular interaction studies for ovine cytochrome B.

**Fig 4:**
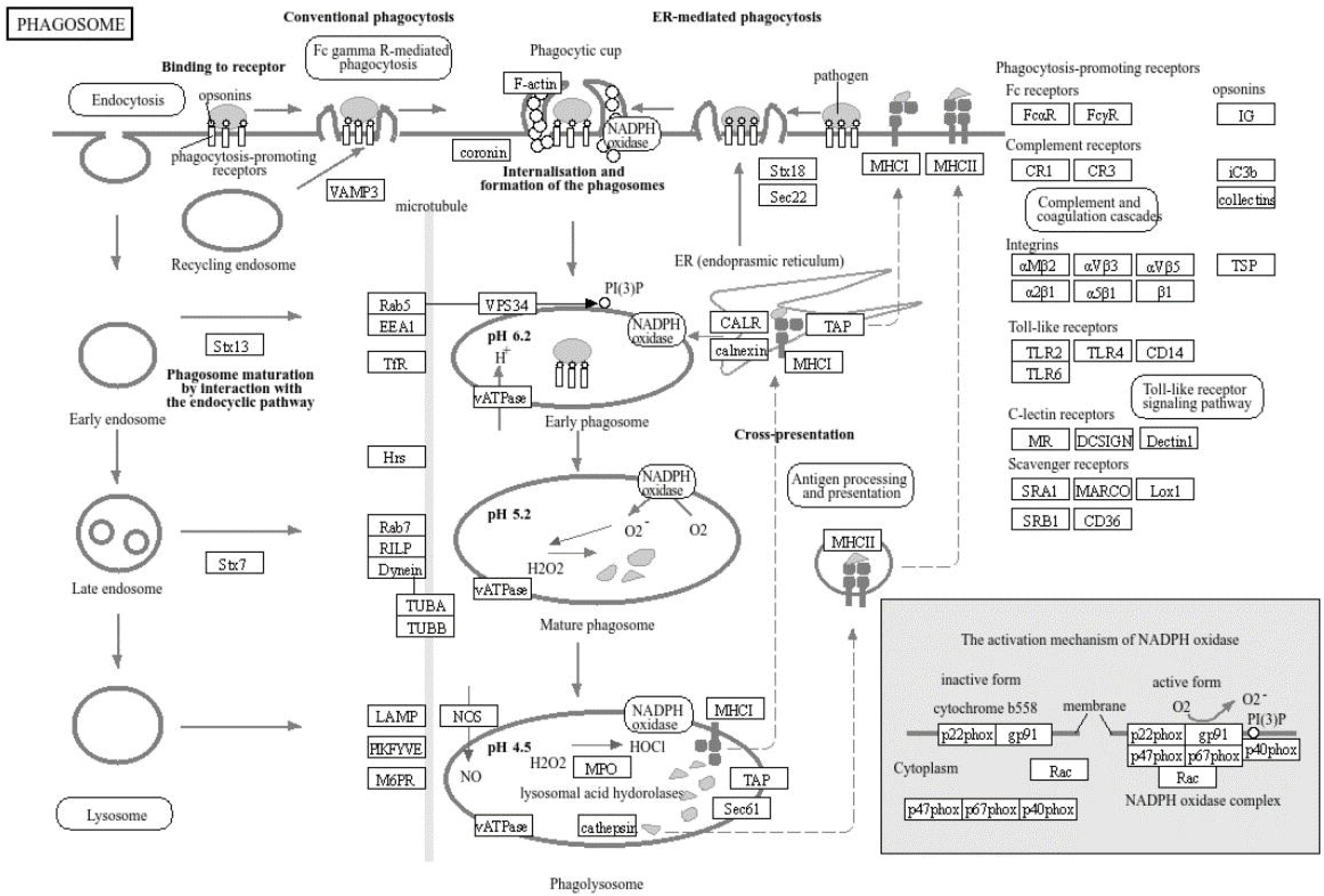
KEGG analysis depicting the role of cytochrome B phagosome pathway.

**Fig 5:**
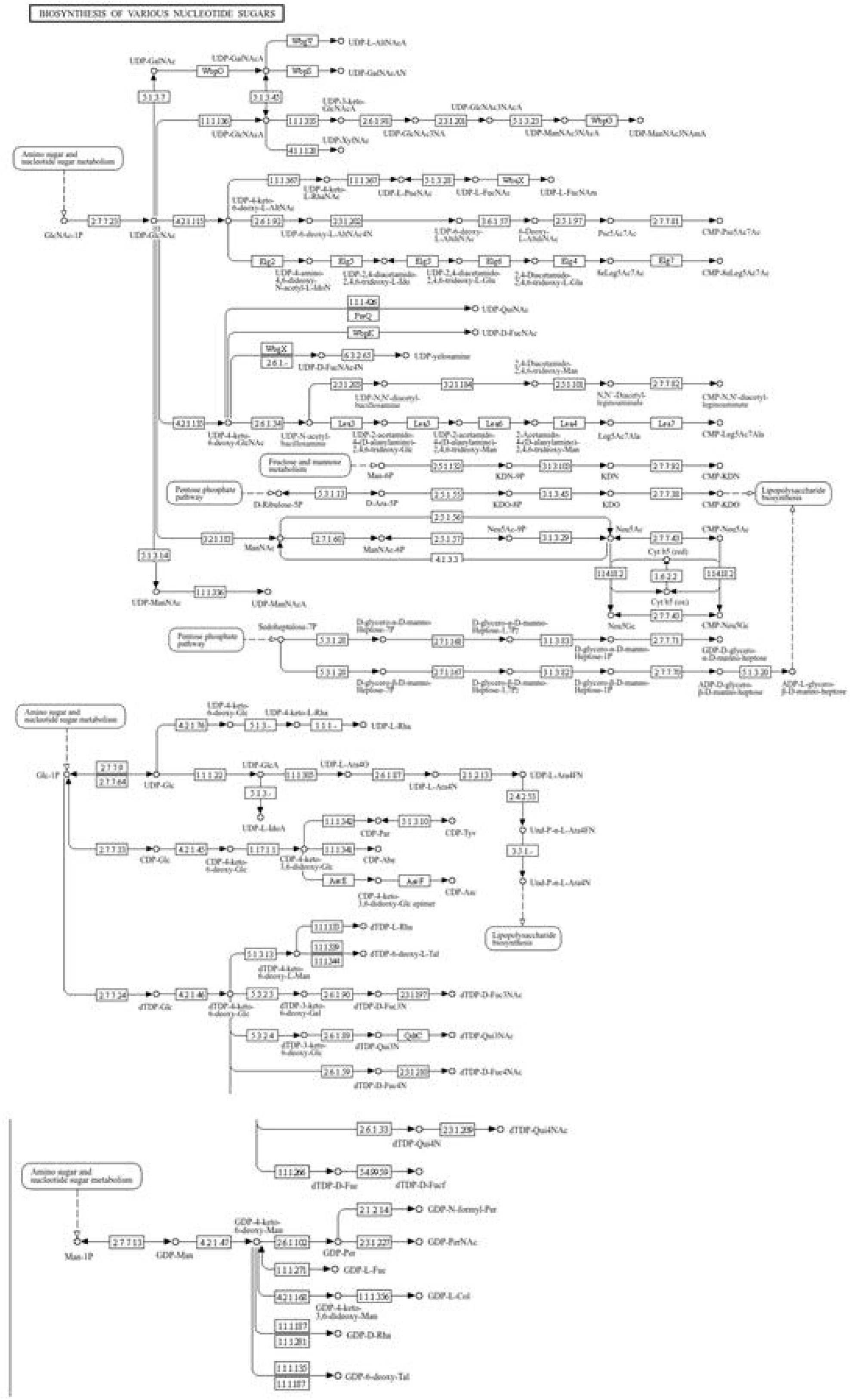
KEGG analysis depicting biosynthesis of various nucleotide sugars.

### Diagnosis for Estimation of faecal egg count for H, contortus in sheep

Strongyle eggs from faeces were visually counted in Macmaster slide under microscope with monitor. We also confirmed that the strongyle eggs were from Haemonchus contortus through presence of adult Haemonchus from abomasum.

### Confirmation of faecal egg/larvae from H. contortus-molecular detection

QPCR is one of the most sensitive technique. The positive samples reflect expression of Cox1 gene for *Haemonchus cortortus* samples in contrast to negative samples.

### Assessment of Haematological and Biochemical parameters of infected sheep in comparison to healthy sheep

We have estimated the haematological and biochemical parameters for the assessment of haematology (Table 5) as well as liver (Table 6) and kidney function test (Table 7). Haemonchus contortus reveals affecting multifunctional. Infected sheep were observed to have significantly reduced haemoglobin concentration (Fig 7) and Total erythrocyte count (Fig 8).

**Fig. 6.**
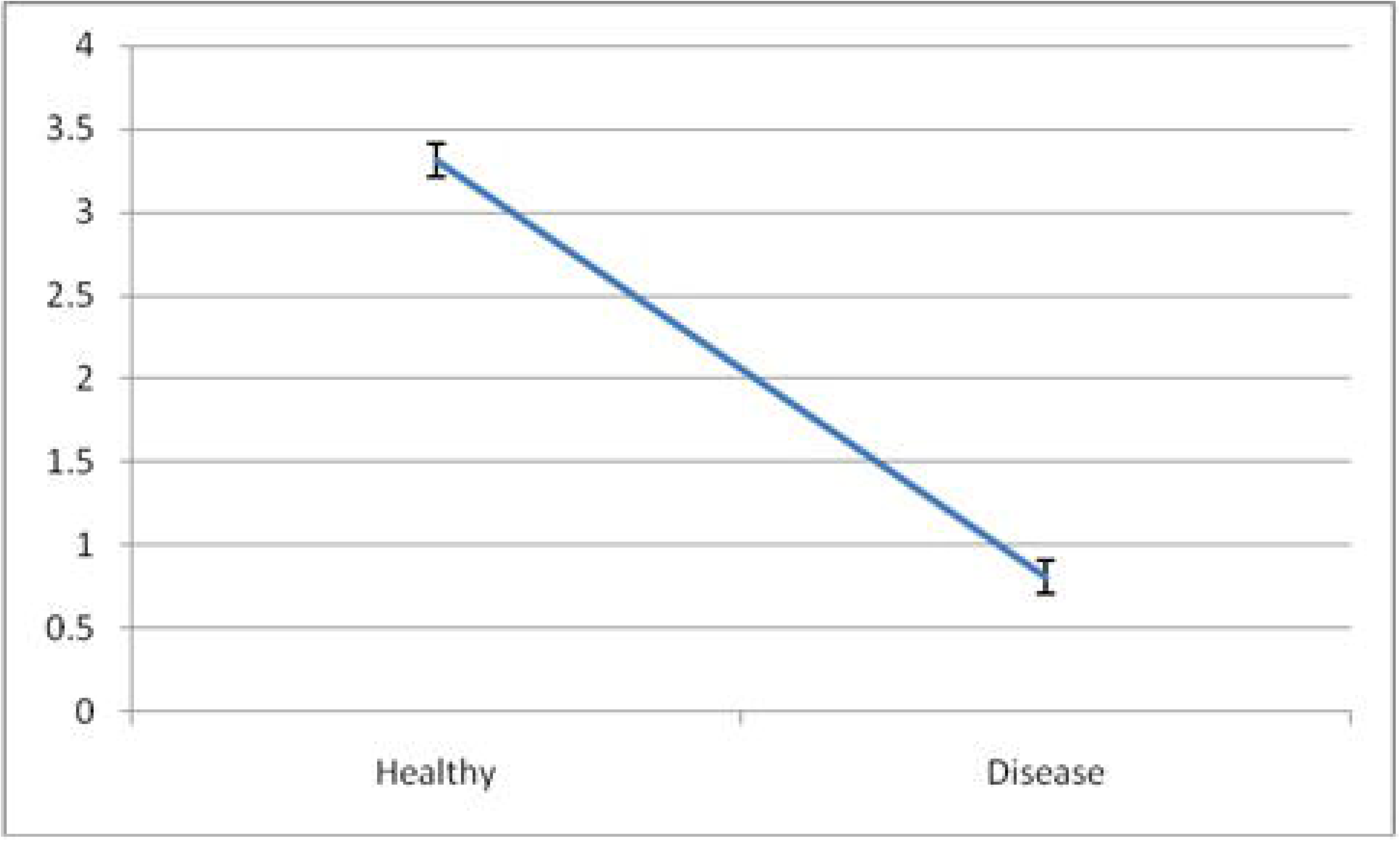
Mean Haemoglobin concentration (gm/dL) in Healthy and infected Sheep w.r.t.H. contortus.

**Fig. 7:**
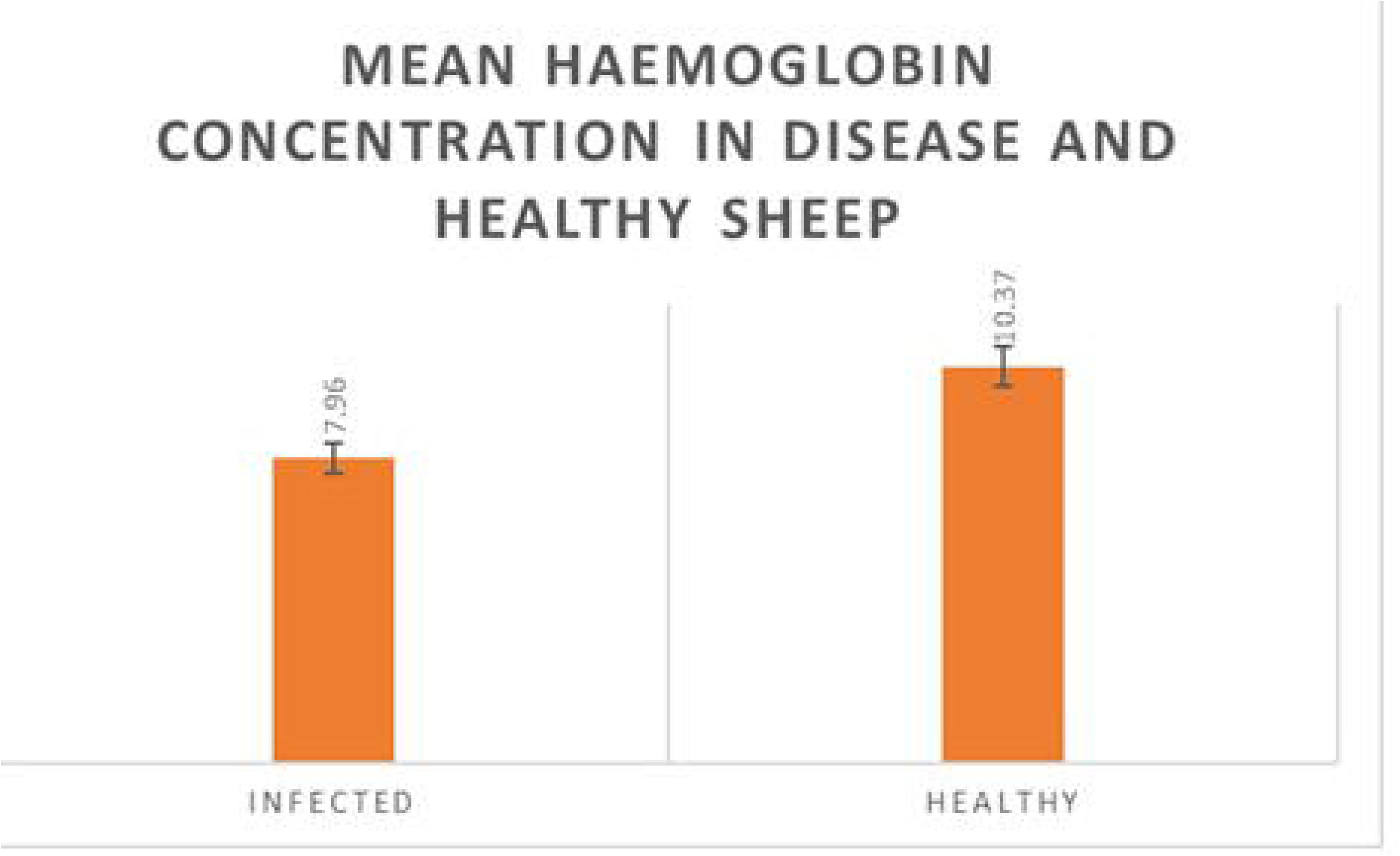
Mean TEC /cumm in Healthy and infected Sheep w.r.t. H.contortus.

**Fig 8:**
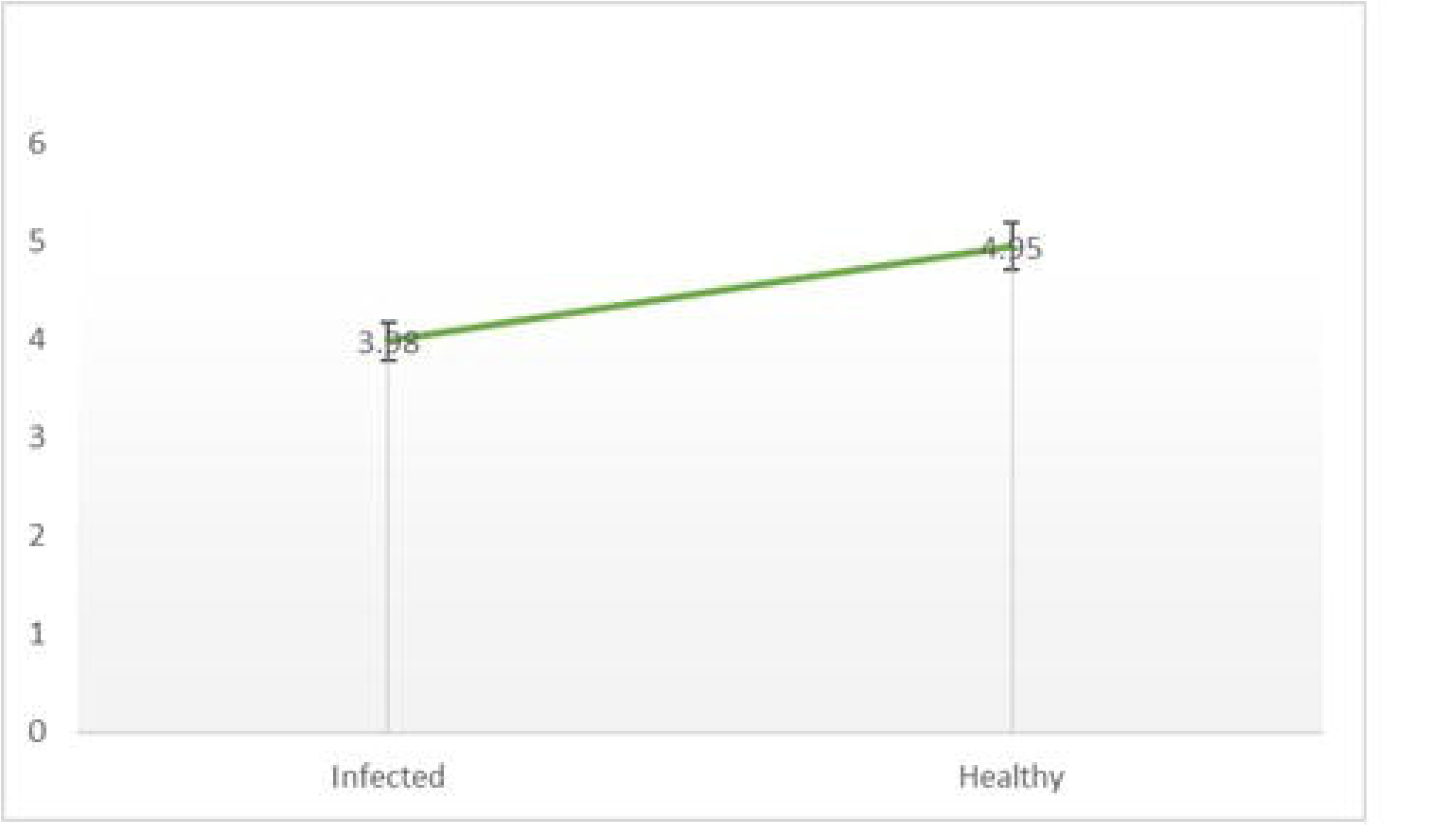
Differential mRNA expression profiling for sheep Cytochrome B with respect to healthy vs. infected ones.

**Table 3:**
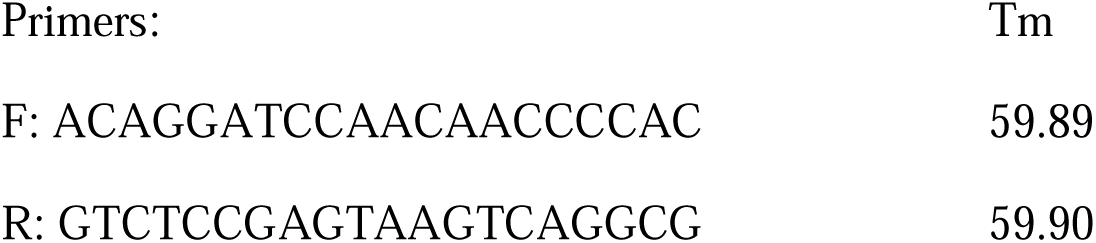
Primers for QPCR analysis of cytochrome B in sheep Primers:

**Table 4:** Estimation of faecal egg count from healthy and infected /diseased sheep.

| Health Status of the sheep | Mean Fecal Egg Count |
| --- | --- |
| Healthy | 50 ± 5.56 |
| Disease | 550 ± 9.45 |

Superscript a and b represent significant differences at P≤0.05

**Table 5:** Haematological parameters for healthy vs. infected sheep.

| S.no | Parameter | Disease | Healthy |
| --- | --- | --- | --- |
| 1. | Hb gm/dl | 7.96<sup>b</sup>±0.19 | 10.37<sup>a</sup>±0.19 |
| 2. | ESR mm | 1.00±0.00 | .90±0.04 |
| 3. | PCV% | 27.84<sup>b</sup>±0.67 | 36.25<sup>a</sup>±0.64 |
| 4. | TEC /cumm | 3.98<sup>b</sup>±0.05 x10<sup>6</sup> | 4.95<sup>a</sup>±0.08x10<sup>6</sup> |
| 5. | TLC/cumm | 9200.00<sup>a</sup>±159.23 | 6587.50<sup>b</sup>±210.81 |
| 5. | Neutrophil % | 53.50<sup>a</sup>±1.00 | 30.25<sup>b</sup>±0.59 |
| 6. | Eosonophil% | 16. 62<sup>a</sup>±0.49 | 2.12<sup>b</sup> ±0.44 |
| 7. | Basophil% | 0.00 | 0.00 |
| 8. | Lymphocyte% | 29.12<sup>b</sup>±0.29 | 60.87<sup>a</sup>±0.29 |
| 9. | Monocyte% | 1.37±0.18 | 1.87±0.29 |

The values that have superscript, was showing significant difference between groups. The superscript 'a' was showing high mean value than 'b'.Significant at ( $p \leq 0.05$ ).

**Table 6:** Biochemical parameters for kidney function test for healthy vs. infected sheep.

| Sr. no | Parameters | Disease | Healthy |
| --- | --- | --- | --- |
| 1 | Total protein (g/dl) | 6.11<sup>b</sup>±0.02 | 7.77<sup>a</sup>±0.03 |
| 2 | Albumin (g/dl) | 1.92<sup>b</sup>±0.04 | 2.80<sup>a</sup>±0.03 |
| 3 | Globulin(g/dl) | 4.18<sup>a</sup>±0.06 | 4.97<sup>b</sup>±0.06 |
| 4 | Albumin:Globulin | 0.46<sup>b</sup>±0.01 | 0.58<sup>a</sup>±0.01 |
| 5. | S.G.P.T IU/L | 39.75<sup>a</sup>±1.4 | 32.12<sup>b</sup>±1.0 |
| 6 | S.G.O.T IU/L | 86.87<sup>a</sup>±2.6 | 68.87<sup>b</sup>±2.7 |
| 7 | Alkaline Phosphatase IU/L | 85.25<sup>a</sup>±3.89 | 64.87<sup>b</sup>±2.09 |
| 8 | Total Bilirubin mg/dl | 0.67<sup>a</sup>±0.07 | 0.37<sup>b</sup>±0.03 |
| 9 | Direct Bilirubin mg/dl | 0.30<sup>a</sup>±0.02 | 0.16<sup>b</sup>±0.01 |
| 10 | Indirect Bilirubin mg/dl | 0.37<sup>a</sup>±0.09014 | 0.21<sup>b</sup>±0.05 |

The Mean values that have superscript, was showing significant difference between groups. The superscript ‘
a’ was showing high mean value than ‘b’.Significant at ( $p \leq 0.05$ ).

**Table 7:**
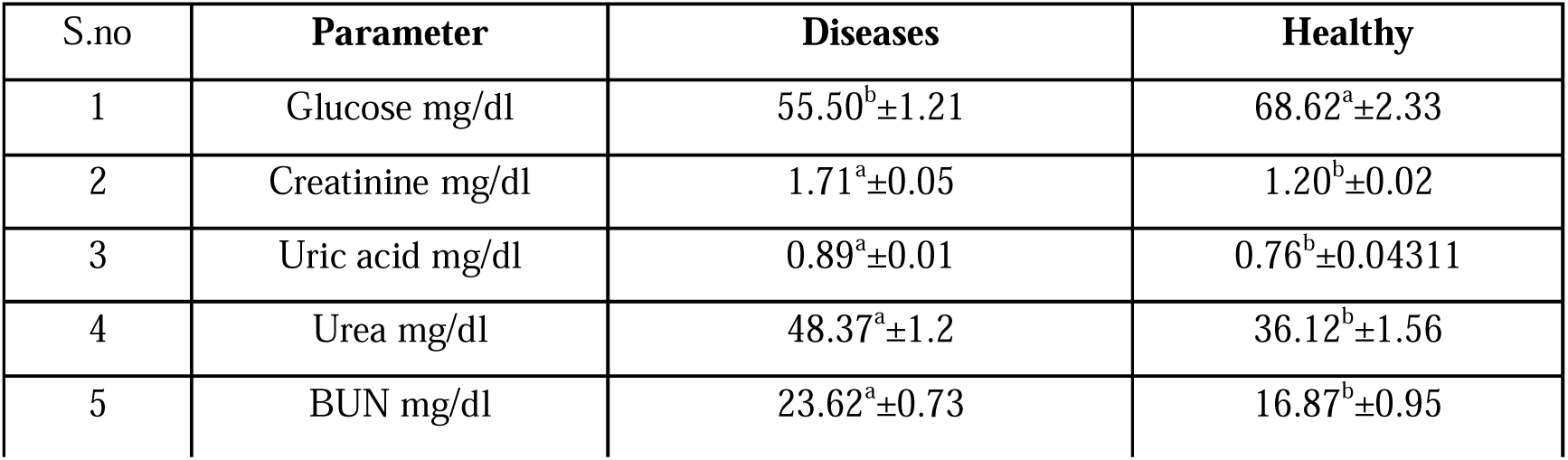
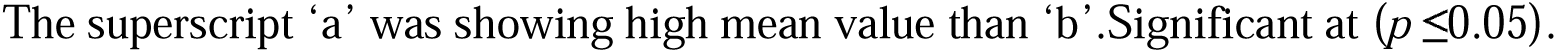
Biochemical parameters for kidney function test for healthy vs. infected sheep. The Mean values that have superscript, was showing significant difference between groups.

| S.no | Parameter | Diseases | Healthy |
| --- | --- | --- | --- |
| 1 | Glucose mg/dl | 55.50<sup>b</sup>±1.21 | 68.62<sup>a</sup>±2.33 |
| 2 | Creatinine mg/dl | 1.71<sup>a</sup>±0.05 | 1.20<sup>b</sup>±0.02 |
| 3 | Uric acid mg/dl | 0.89<sup>a</sup>±0.01 | 0.76<sup>b</sup>±0.04311 |
| 4 | Urea mg/dl | 48.37<sup>a</sup>±1.2 | 36.12<sup>b</sup>±1.56 |
| 5 | BUN mg/dl | 23.62<sup>a</sup>±0.73 | 16.87<sup>b</sup>±0.95 |

The superscript 'a' was showing high mean value than 'b'. Significant at ( $p \leq 0.05$ ).

### Differential mRNA expression profiling for Cytochrome B gene from abomassum of Sheep

We have analyzed the abomassum of sheep from both infected and healthy counterpart. We observed better expression profile for Cyt B in healthy sheep in comparison to infected ones (Fig8).

## Discussion

Despite significant advances in parasite immunology research, challenges remain in understanding how precisely helminths interact with their hosts and tolerate the complex immune responses generated against them, and how to utilize such information to define new ways of combating these parasitic infections. If prophylactic vaccines or immunotherapy for helminths are to be developed as an alternative to anthelmintics, it will be essential to better understand the immune mechanisms that provide protection against these helminth infections. The most promising recent development in this field has been the ongoing identification and characterization of molecules that are involved in the interaction between helminths and immune cells of their hosts.

In our current study, we observe better expression of cytochrome B, indicative of better mitochondrial DNA copy numbers in healthy sheep in comparison to sheep infected with Haemonchus contortus. We had diagnosed the infected sheep through symptoms, faecal egg count, visualization of adults in abomasum and finally through molecular detection with QPCR with Cox1 gene primer from *Haemonchus contortus*. In our lab, we earlier observed better expression of RIGI^17^, CD14^18^, IL6^20^, IL10^20^ in healthy sheep in comparison to infected ones. There exists certain mechanism how mitochondria can function in immune response. Mitochondria can release mitochondrial DNA (mtDNA) and other mitochondrial components in response to infection or cellular stress. This extracellular mtDNA acts as a danger-associated molecular pattern (DAMP), triggering the innate immune system’s response, including the release of pro-inflammatory cytokines and the activation of various immune cells^42^. Mitochondria acts through a complex mechanism of interaction of nuclear genes^9^.

We also observed molecular interaction of cytochrome B gene with other genes, most of them are of mitochondrial origin along with some nuclear origin ubiquinone genes. We detected four iron binding sites, since cytochrome B is equally involved with heme formation. *Haemonchus contortus* is a gastrointestinal parasite affect the gut of sheep, abomasum particularly leading to heavy blood loss. Mortality is mostly recorded in young ones with lesser blood volume and body size. Cytochrome B is directly linked to heme production and iron binding, so indirectly helps in hematopoiesis. As a result, young lambs with lesser copy number of mitochondrial DNA, manifested as lesser expression of cytochrome B will have less blood production and volume, so subjected to higher mortality with *Haemonchus contortus* infection. Cytochrome is also responsible for apoptosis as revealed in ubiqinone binding site, that may have indirect effect on immune response for sheep. We have observed through KEGG analysis pathway that CytB is equally involved with phagosome formation, which may aid in engulfing of parasites. They are equally responsible for biosynthesis of nucleotide sugar, evident from biochemical pathway analysis which aid in DNA and RNA formation and ultimately aid in immune response.

Mitochondria has been playing a multifaceted role in oxidative phosphorylation, calcium metabolism, iron metabolism and immune functions against pathogens, particularly against bacteria^4,9^, and virus^43^. Mitochondria play a crucial role in the innate immune response, which is the first line of defense against parasites. In our earlier studies, we have detected certain mutations in cytochrome B gene in sheep causing extreme debility in sheep with cardiac and other physiological involvement^1^.

Mitochondria aid in Metabolic Signaling. Mitochondria are central to cellular metabolism, and their dysfunction can impact the host’s ability to mount an effective immune response. The host’s ability to produce energy, generate reactive oxygen species (ROS), and maintain proper cellular redox balance all rely on mitochondrial function, which, in turn, affects immune cell function and parasite killing^44^.Any defect in mitochondrial metabolic signalling leads to lowered immune function and diseases.

Mitochondria play a central role in apoptosis, a programmed cell death process. When cells are infected with parasites, they can initiate apoptosis as a defense mechanism to prevent the spread of the infection. This process is regulated in part by mitochondrial factors, including the release of cytochrome c from mitochondria, which triggers the apoptotic cascade. Mitochondria plays a significant role in the controlling Inflammation. Mitochondrial dysfunction may cause an increased production of reactive oxygen species (ROS), leading to increase inflammation. Dysregulated inflammation can be harmful in the context of parasite infections, causing collateral tissue damage. Mitochondrial signaling can impact the balance between pro-inflammatory and anti-inflammatory responses^45^. Mitochondrial Antiviral Signaling (MAVS) is a mitochondrial protein that plays a crucial role in the host’s antiviral defense^46^. It activates signaling pathways leading to the production of interferons, which are key molecules in the defense against viral infections. Although this aspect is more related to viral infections, it showcases the involvement of mitochondria in antiparasitic immunity.

In our earlier study, we observed increased expression of RIGI in healthy sheep^17^. RIGI is a pattern recognition receptor acts through MAVS as reported in case of viral infection. Here also we report more mitochondrial copy number in healthy which may be directly or indirectly linked to RIGI in destruction of parasitic pathogen. Through molecular docking studies, we observed, RIGI binds with Haemonchus surface proteins. Such mitochondrial-nuclear cross talk might be responsible for a comprehensive immune response process against dreadly parasite as Haemonchus contortus^17^.

Mitochondrial is functionally versatile, which is mostly performed through changes in its morphology. For example, cytosolic signals can regulate the efficiency of fuel usage by mitochondria by impinging on fission, fusion, and cristae remodeling^47^. During starvation, when autophagy is induced to liberate nutrients and mobilize FAs from LDs that undergo lipophagy, mitochondria elongate and their cristae surface increases, to evade autophagic degradation and increase the efficiency of ATP production^48–52^. Furthermore, these morphological changes enhance the efficiency of b-oxidation by increasing mitochondrial association with and uptake of FAs from LDs^53^. Notably, host cell mitochondria are recruited to the vacuolar membranes that harbor a diverse group of evolutionarily distinct intracellular pathogens, including the bacteria *Legionella pneumophila* and *Chlamydia psittaci*, as well as the protozoan parasite *Toxoplasma gondii*^54^, all of which undergo replication within these compartments. *T. gondii* infects approximately one-third of the global human population and exhibits an exceptionally broad host range across vertebrate species. Similar to *Chlamydia*, it extensively exploits host-derived lipid species to support its intracellular growth and survival.

Morphology of mitochondria in response to infection depends on the microbe studied: mitochondria elongate around the Toxoplasma vacuole^54^; in contrast, they fragment following Listeria entry or Vibrio cholerae infection, a process essential to promote their growth. The mechanism of mitochondrial elongation early around the Toxoplasma vacuole is yet to explore. Using Toxoplasma to study there relationship among an intracelllular microbe, access to essential lipids, and mitochondrial morphology, we show that Toxoplasma co-optshost cell lipophagy to obtain FAs needed for its proliferation. Conversely, host cell mitochondria undergo fusion around the *Toxoplasma* parasitophorous vacuole, thereby sequestering fatty acids and limiting parasite proliferation.

It is quite interesting to note that mitochondria was observed to be absent in some parasites, namely Dinoflagelles^55^, giardia (with rudimentary mitosome). This may be due to their ability to directly consuming energy from the host.

Earlier we have reported the role of RIGI gene to have potent role in parasitic immunity against Haemonchus contortus, where it binds to the parasite and binding sites predicted to be Histidine 302 position to the Proline 763 position for the surface proteins alpha tubulin and beta tubulin. a– and b-tubulin, being major constituents of microtubules, bind two moles of GTP, one at an exchangeable site on the alpha chain and one at a nonexchangeable site on the beta chain.. we identified the binding site for RIG-I with the polypeptide of parasite at 300-750 amino acid for most of the protein. The parasitic proteins (Alpha tubulin, Beta tubulin, lectin, galectin, and cysteine protease) were found to be the most promising binding site with ovine RIG-I. This was the first report for role of RIGI in parasitic infection, earlier only antiviral effect were known. After binding with the pathogen, RIGI acts through MAVS and secretes interleukins that ultimately destroys the pathogen. Our study reported that RIGI binds to *Haemonchus contortus*^17^ and recruit to MAVS (mitochondrial protein), in turn secretes IL6, IL10, which ultimately destroys the pathogen. We have also observed increased expression of IL6 and IL10 in healthy sheep in comparison to the infected ones^20^. Similarly CD14 was also reported as receptor for Haemonchus contortus for the first time)^18^. The structural alignment of alpha tubulin of H. contortus with the CD14 gene, and aligned site being depicted as Glu27 to Ala284. The structural alignment of betatubulin of H.contortus with the CD14 gene, and the aligned site is depicted as Glu27 and Ala284.

Haemoglobin concentration as well as total erythrocyte count was observed to be significantly lower in infected sheep compared to healthy ones. We observed lower mitochondrial copy number as manifested through cytochrome B expression profile. We have already documented four iron binding sites in cytochrome B responsible for haem production and ultimately for haemoglobin synthesis. Better haemoglobin synthesis leads to more production of erythrocyte, manifested as erythrocytic count. Thus we observe infected sheep with lower erythrocyte count as well as lower copy number for mitochondria. Another mode for destruction of parasite by mitochondria is through Phagosome formation, as depicted in molecular pathway analysis. Hence mitochondrial DNA seems to have great influence against parasitic immunity^57^.

In recent days researches are being undertaken for assessment of mitochondrial copy number is association to various pathological conditions^56–67^. Reports indicate that mitochondrial copy number varies from tissue to tissue^60,66^ and also species wise variation exists^62^. Various methodologies for estimation of mitochondrial copy number have been elucidated as QPCR based differential mRNA expression profile relative to nuclear DNA quantity, Digital PCR^63^, DNA sequencing^64^. Mitochondrial copy number has been associated to Personality traits^57^, Low-Density Lipoprotein Cholesterol^58^, and Cardiovascular Disease Risk, ^58^ metastatic breast cancer^59^, Multiple Sclerosis^62^ **and** metabolic traits^64^.

## Conclusion

Mitochondria through its copy number variation plays an immense role in parasitic immunity against *Haemonchus contortus* in sheep model. We have characterized cytochrome B from gut of sheep and observed four iron binding sites, responsible for heme production leading to better haemoglobin as well as total erythrocytic count for healthy sheep with more mitochondrial DNA copy numbers, in comparison to diseased ones. Another possible mode for destruction of parasite by mitochondria is through phagosome production, and biosynthesis of various nucleotide sugar. Additionally, RIGI binds to surface protein alpha tubulin and beta tubulin of the *Haemonchus contortus,* which is then recruited to *mitochondrial* protein MAVS, releasing IL6, IL10.

## Acknowledgement

The authors are thankful to Department of Biotechnology, Ministry of Science and Technology, Govt. of India (Grant number BT/Bio-CARe/04/10100/2013-14). The funders had no role in study design, data collection and analysis, decision to publish, or preparation of the manuscript. The authors are equally thankful to Vice-Chancellor, West Bengal University of Animal and Fishery Sciences for providing the financial support. Thanks to Director, AH & VS, Animal Resource Development Department, Govt. of West Bengal.

## Declaration

This is to declare that the study has been approved by Institutional Biosafety Committee, West Bengal University of Animal and Fishery Sciences.

## References

1. Pal, A., Pal, Ab., Banerjee, S., Batobyal, S. and Chatterjee, P.N. 2019. Mutation in *Cytochrome B gene* causes debility and adverse effects on *health* of sheep. Mitochondrion 46: 393–404.10.1016/j.mito.2018.10.003

2. Pradhan, M., Pal, A., Samanta, A.K., Banerjee, S., Samanta, R. 2018. Mutations in *cytochrome B gene* effects female reproduction of Ghungroo pig. Theriogenology. 119: 121–130. doi: 10.1016/j.theriogenology.2018.05.015

3. Farge G, Falkenberg M. Organization of DNA in Mammalian Mitochondria. Int J Mol Sci. 2019 Jun 5;20(11):2770. doi: 10.3390/ijms20112770. PMID: 31195723; PMCID: PMC6600607.

4. Sahu, J., Pal, A et al., 2023. Role of NUMTs (Nuclear mitochondrial DNA) genes in affecting disease resistance in duck against Pasteurellosis. bioRxiv. doi: 10.1101/2023.08.30.555528;

5. Lee SR, Han J. Mitochondrial Nucleoid: Shield and Switch of the Mitochondrial Genome. Oxid Med Cell Longev. 2017;2017:8060949. doi: 10.1155/2017/8060949.

6. Falkenberg M, Larsson NG, Gustafsson CM. Replication and Transcription of Human Mitochondrial DNA. Annu Rev Biochem. 2024 Aug;93(1):47–77. doi: 10.1146/annurev-biochem-052621-092014.

7. J. Lu, L.K. Sharma, Y. Bai, Implications of mitochondrial DNA mutations and mitochondrial dysfunction in tumorigenesis, Cell Res. 19 (2009) 802–815.

8. F.M. Yakes, B. Van Houten, Mitochondrial DNA damage is more extensive and persists longer than nuclear DNA damage in human cells following oxidative stress, Proc. Natl. Acad. Sci. U. S. A. 94 (1997) 514–519.

9. Chakraborty, A., Pal, A. 2023. Mitochondrial genes affect the immune response against bacterial infection (Duck Pasteurellosis) through a cascade of mechanisms mediated by nuclear immune genes-a cross talk with nuclear and mitochondrial gene. bioRxiv preprint doi: 10.1101/2023.08.30.555483

10. Pal, A. 2021. Protocols in Advanced Genomics and allied techniques., Springer Publication. Springer Protocols Handbooks. ISBN 978-1-0716-1817-2 ISBN 978-1-0716-1818-9 (eBook),10.1007/978-1-0716-1818-9

11. Pal, A. and Chakravarty, A.K. 2019. Genetics & Breeding for the Disease resistance of livestock. Elsevier Publication. Academic publisher. ISBN: 9780128164068

12. Peter, J. W., and Chandrawathani, P. (2005). Haemonchus contortus: parasite problem No. 1 from tropics-polar Circle. Problems and prospects for control based on epidemiology. Trop. Biomed. 22 (2), 131–137.

13. Pal, A. 2018. Birbhum sheep-the prolific sheep of West Bengal, Pp: 51.Published by DREF. West Bengal University of Animal and Fishery Sciences, West Bengal, India. ISBN: 978-93-5268-187-7

14. Pal, A., Banerjee, S., Karmakar P., 2022. Phylophenomic and Phylogenomic analysis for Ovis aries reveals distinct identity of newly reported breed. BioRxiv, 2022.07. 31.502249

15. Fissiha W, Kinde MZ. Anthelmintic Resistance and Its Mechanism: A Review. Infect Drug Resist. 2021 Dec 15;14:5403–5410. doi: 10.2147/IDR.S332378.

16. J. Charlier, D.J. Bartley, S. Sotiraki, M. Martinez-Valladares, E. Claerebout, G. von Samson-Himmelstjerna, S.M. Thamsborg, H. Hoste, E.R. Morgan, L. Rinaldi, Chapter Three – Anthelmintic resistance in ruminants: challenges and solutions, Editor(s): David Rollinson, Russell Stothard, Advances in Parasitology, Academic Press, Volume 115, 2022, Pages 171–227, ISBN 9780323988711, 10.1016/bs.apar.2021.12.002.

17. Banerjee S, Pal, A., Mandal, S.C., Chatterjee, P. N., Chatterjee, J.K. 2021. RIG-I has a role in *immunit*y against *Haemonchus contortus,* a gastrointestinal parasite in *Ovis aries-* a novel report. Frontiers in immunology. 08 January 2021 10.3389/fimmu.2020.534705.

18. Rawat, K., Pal, A., Banerjee, S., Pal, Ab., Mandal, S.C, Batabyal, S. 2021. Ovine CD14-an immune response gene has a role against gastrointesinal nematode *Haemonchus contortus* –a novel report. Frontiers in Immunology. 10.3389/fimmu.2021.664877

19. Banerjee, S., Pal, A., Pal, Ab., Chatterjee, J.K. 2023. Ovine MyD88 have active role in host resistance against Haemonchus contortus infection. doi: 10.1101/2023.10.08.561398

20. Rawat, K., Pal, A et al., 2023. Role of IL-6 and IL10 immune response genes against nematode Haemonchus contortus in ovis aries-a novel report. bioRxiv preprint doi: 10.1101/2023.08.30.555538

21. Pal, A., Banerjee, S., Pathak, K., Das, M.K., Chatterjee, P.N. 2022. Molecular phylogenetic analysis for Ovis aries with whole mitochondrial genome sequencing. doi. 10.1101/2022.07.29.502105

22. Pal, A., Chakraborty, A., Debnath, M. 2022. Whole mitochondrial genome sequencing-a novel approach for studying Phylogenomics for Anas platyrynchos. doi: 10.1101/2022.07.11.499548

23. Pal, A., Chakraborty, A., Debnath, M. 2022. Molecular evolution and characterization of domestic duck (Anas platyrynchos) and Goose (Anser indicus) with reference to its wild relatives through whole mitochondrial genome sequencing. doi: 10.1101/2022.07.19.500621

24. Benjamin MM. In: Outline of Veterinary Clinical Pathology. 3rd Edn. New Delhi: Kalyani Pub. (1985) p. 233–54.

25. Hawk PB. In: Hawak’s Physiological Chemistry. 14th Edn. London: McGraw Hill Book Co (1965). 218 p.

26. Jain NC. In: Essentials of Veterinary Haematology. Pennsylvania: Lea and Febiger Publishers Malvern (1986) p. 57–62.

27. Murmu, A.K., Pal, A., et al., 2023. Role of mucin gene for growth in Anas platyrynchos – a novel report. Frontiers Veterinary science. doi I 10.3389/fvets.2023.1089451

28. Pal, A, Pal, A. Jr., Mallick, A.I., Biswas, P. and Chatterjee, P.N. 2019. Molecular characterization of *Bu-1 and TLR2* gene in Haringhata Black chicken. Genomics 112(1): 472–483. 10.1016/j.ygeno.2019.03.010

29. Pal, A., Pal, Ab., Baviskar, P. 2021. Molecular characterization of RIGI, TLR7 and TLR3 as immune response gene of indigenous ducks in response to Avian influenza. doi: 10.3389/fmolb.2021.633283 (Frontier Molecular BioScience)

30. Pal, S., Pal, A., 2022. Role of RIGI, MDA5 and interferon alpha of duck in Duck Plague infection – a novel report. doi: 10.1101/2022.01.26.477779

31. Debnath, M., Pal, A., Chakraborty, A., Pal, S., Pal, Ab. 2022. Genomics for reproduction in Anas platyrynchos-a novel report. doi: 10.1101/2022.05.29.493861

32. Glick DM, et al. Glossary of Biochemistry and Molecular Biology, Portland Press, London, UK, Revised edition(1977).

33. Petersen B, et al.A generic method for assignment of reliability scores applied to solvent accessibility predictions. BMC Structural Biology, 9, 51. doi: 10.1186/1472-6807-9-51(2009).

34. Ebina T, et al.Loop-length dependent SVM prediction of domain linkers for high-throughput structural proteomics. Biopolymers 92(1), 1–8(2009).

35. Cunningham MC, et al.CD14 Employs Hydrophilic Regions to “Capture” Lipopolysaccharides. Journal of Immunology 164, 3255–3263(2000).

36. Muroi M, et al. Regions of the mouse CD14 molecule required for toll-like receptor 2– and 4-mediated activation of NF-kappa B. Journal of Biological Chemistry; 277(44), 42372–42379(2002).

37. Kiefer F, Arnold K, Künzli M, Bordoli L, Schwede T. The SWISS-MODEL Repository and associated resources. Nucleic Acids Res. 2009 Jan;37(Database issue):D387–92. doi: 10.1093/nar/gkn750.

38. Kelley L. et al.The Phyre2 web portal for protein modeling, prediction and analysis. Nature Protocols; 10,845–854. doi: 10.1038/nprot.2015.053(2015).

39. Wiederstein M., et al. ProSA-web: interactive web service for the recognition of errors in three-dimensional structures of proteins, Nucleic Acids Research, Volume 35, Issue suppl_2, Pages W407–W410, 10.1093/nar/gkm290(2007)

40. Zhang Y, et al. TM-align: A protein structure alignment algorithm based on TM-score, Nucleic Acids Research, 33: 2302–2309(2005).

41. Franceschini A, Szklarczyk D, Frankild S, Kuhn M, Simonovic M, Roth A, Lin J, Minguez P, Bork P, Von Mering C, Jensen LJ. STRING v9. 1: protein-protein interaction networks, with increased coverage and integration. Nucleic acids research (2012) 780 41(D1):D808–15. doi:10.1093/nar/gks1094

42. Kim, J., Kim, HS. & Chung, J.H. Molecular mechanisms of mitochondrial DNA release and activation of the cGAS-STING pathway. Exp Mol Med 55, 510–519 (2023). 10.1038/s12276-023-00965-7

43. Duan X, Liu R, Lan W, Liu S. The Essential Role of Mitochondrial Dynamics in Viral Infections. Int J Mol Sci. 2025 Feb 24;26(5):1955. doi: 10.3390/ijms26051955.

44. Yang S, Lian G. ROS and diseases: role in metabolism and energy supply. Mol Cell Biochem. 2020 Apr;467(1-2):1–12. doi: 10.1007/s11010-019-03667-9. Epub 2019 Dec 7. Erratum in: Mol Cell Biochem. 2020 Apr;467(1-2):13. doi: 10.1007/s11010-020-03697-8.

45. Marchi, S., Guilbaud, E., Tait, S.W.G., et al. Mitochondrial control of inflammation. Nat Rev Immunol 23, 159–173 (2023). 10.1038/s41577-022-00760

46. Ren Z, Ding T, Zuo Z, Xu Z, Deng J, Wei Z. Regulation of MAVS Expression and Signaling Function in the Antiviral Innate Immune Response. Front Immunol. 2020 May 27;11:1030. doi: 10.3389/fimmu.2020.01030.

47. Pernas, L. and Luca Scorrano, L. 2016. Mito-Morphosis: Mitochondrial Fusion, Fission, and Cristae Remodeling as Key Mediators of Cellular Function Annual Review of Physiology, vol 78 10.1146/annurev-physiol-021115-105011.

48. Gomes, L.C., Di Benedetto, G., Scorrano, L. During autophagy mitochondria elongate, are spared from degradation and sustain cell viability *Nat*. Cell Biol. 2011; 13:589–598 DOI: 10.1038/ncb2220

49. Patten, D.A., Wong, J., Khacho, M. OPA1-dependent cristae modulation is essential for cellular adaptation to metabolic demand. EMBO J. 2014; 33:2676–2691 DOI: 10.15252/embj.201488349

50. Rambold, A.S., Kostelecky, B., Elia, N. Tubular network formation protects mitochondria from autophagosomal degradation during nutrient starvation *Proc*. Natl. Acad. Sci. USA. 2011; 108:10190–10195. DOI: 10.1073/pnas.1107402108

51. Reggiori, F., Klionsky, D.J. Autophagy in the eukaryotic cell. Eukaryot. Cell. 2002; 1:11–21. DOI: 10.1128/EC.01.1.11-21.2002

52. Singh, R., Kaushik, S., Wang, Y. Autophagy regulates lipid metabolism Nature. 2009; 458:1131–1135. doi: 10.1038/nature07976.

53. Herms A, Bosch M, Reddy BJ, Schieber NL, Fajardo A, Rupérez C, Fernández-Vidal A, Ferguson C, Rentero C, Tebar F, Enrich C, Parton RG, Gross SP, Pol A. AMPK activation promotes lipid droplet dispersion on detyrosinated microtubules to increase mitochondrial fatty acid oxidation. Nat Commun. 2015 May 27;6:7176. doi: 10.1038/ncomms8176.

54. Lena Pernas, Camilla Bean, John C. Boothroyd, and Luca Scorrano. 2018. Mitochondria Restrict Growth of the Intracellular Parasite Toxoplasma gondii by Limiting Its Uptake of Fatty Acids Cell Metabolism 27, 886–897

55. Kayal E, Smith DR. Is the Dinoflagellate Amoebophrya Really Missing an mtDNA? Mol Biol Evol. 2021 May 19;38(6):2493–2496. doi: 10.1093/molbev/msab041.

56. Xing, et al., 2008. Mitochondrial DNA Content: Its Genetic Heritability and Association With Renal Cell Carcinoma. J Natl Cancer Inst 2008;100: 1104–1112

57. Oppong RF, Terracciano A, Picard M, Qian Y, Butler TJ, Tanaka T, Moore AZ, Simonsick EM, Opsahl-Ong K, Coletta C, Sutin AR, Gorospe M, Resnick SM, Cucca F, Scholz SW, Traynor BJ, Schlessinger D, Ferrucci L, Ding J. Personality traits are consistently associated with blood mitochondrial DNA copy number estimated from genome sequences in two genetic cohort studies. Elife. 2022 Dec 20;11:e77806. doi: 10.7554/eLife.77806.

58. Liu, et al, 2023. Association Between Whole Blood–Derived Mitochondrial DNA Copy Number, Low-Density Lipoprotein Cholesterol, and Cardiovascular Disease Risk. J Am Heart Assoc. 2023;12:e029090. DOI: 10.1161/JAHA.122.029090

59. Rai, N.K., Panjwani, G., Ghosh, A.K., Haque, R., Sharma, L.K. 2021, Analysis of mitochondrial DNA copy number variation in blood and tissue samples of metastatic breast cancer patients (A pilot study). Biochemistry and Biophysics Reports 26 (2021) 100931.

60. Naue J, Xavier C, Hörer S, Parson W, Lutz-Bonengel S. Assessment of mitochondrial DNA copy number variation relative to nuclear DNA quantity between different tissues. Mitochondrion. 2024 Jan;74:101823. doi: 10.1016/j.mito.2023.11.006.

61. Longchamps RJ, Castellani CA, Yang SY, Newcomb CE, Sumpter JA, Lane J, et al. (2020) Evaluation of mitochondrial DNA copy number estimation techniques. PLoS ONE 15(1): e0228166. 10.1371/journal.pone.0228166.

62. Leuthner TC, Hartman JH, Ryde IT, Meyer JN. PCR-Based Determination of Mitochondrial DNA Copy Number in Multiple Species. Methods Mol Biol. 2021;2310:91–111. doi: 10.1007/978-1-0716-1433-4_8.

63. Wendy K. Shoop, Cassandra L. Gorsuch, Sandra R. Bacman, Carlos T. Moraes.2022.Precise and simultaneous quantification of mitochondrial DNA heteroplasmy and copy number by digital PCR, Journal of Biological Chemistry, 298(11), 2022, 10.1016/j.jbc.2022.102574.

64. Ganel L, Chen L, Christ R, Vangipurapu J, Young E, Das I, Kanchi K, Larson D, Regier A, Abel H, Kang CJ, Scott A, Havulinna A, Chiang CWK, Service S, Freimer N, Palotie A, Ripatti S, Kuusisto J, Boehnke M, Laakso M, Locke A, Stitziel NO, Hall IM. Mitochondrial genome copy number measured by DNA sequencing in human blood is strongly associated with metabolic traits via cell-type composition differences. Hum Genomics. 2021 Jun 7;15(1):34. doi: 10.1186/s40246-021-00335-2.

65. Sabaie, H., Taghavi Rad, A., Shabestari, M., et al. Mitochondrial DNA Copy Number as a Hidden Player in the Progression of Multiple Sclerosis: A Bidirectional Two-Sample Mendelian Randomization Study. Mol Neurobiol 62, 11643–11653 (2025). 10.1007/s12035-025-04980-9

66. Rath, S.P., Gupta, R., Todres, E., Wang, H., Jourdain, A.A., Ardlie, K.G., Calvo, S.E., Mootha, V.K. 2024. Mitochondrial genome copy number variation across tissues in mice and humans. NAS 2024, 121 (33).10.1073/pnas.2402291121

